# Effect of Sulfate Availability on Phytoplankton Stoichiometry

**DOI:** 10.1101/2024.05.04.589031

**Authors:** Larisa Whitney, Mariona Segura-Noguera, Zuoxi Ruan, Mario Giordano

## Abstract

Sulfur is a key element in multiple metabolic pathways of phytoplankton cells. The effect of environmental S availability on phytoplankton elemental quotas and stoichiometry has been addressed in few studies, using a limited number of species and with contradictory results.
Using high-temperature combustion oxidation and X-ray fluorescence methods, we measured the concentrations of micro- and trace elements in monocultures of 20 phytoplankton species, grown with different sulfate concentrations representing those of early and modern oceans.
The red lineage species, with higher S requirement and metabolic S fluxes, have higher S content than those of the green lineage, resulting in lower C:S (93) and higher S:P (1.06) than the green lineage species (226 and 0.76, respectively). Zn was the only trace element affected by the environmental concentration of sulfate.
Phytoplankton cells respond to different sulfate availability by either increasing Zn or decreasing P quotas, where the P response is more characteristic in the red lineage, and the Zn response is independent of genotypic constraints or plastid type. This study sheds light on a metabolic basis for the expansion of the red lineage algae and their current dominance in ocean waters.

**Plain language summary:** Microalgae assimilate dissolved sulfate as a source of sulfur, an element that takes part in multiple biochemical pathways and cellular functions. Changes in the concentration of dissolved sulfate in the environment have an effect on the cellular dynamics of several essential biological elements, essencially phosphorus and zinc. The results of this study help to understand the biogeochemical cycling of these elements in the ocean and the dominance of certain phytoplankton lineages.

## Introduction

The debate on the existence and extent of a generalized elemental stoichiometry shared among phytoplankton remains an active focus within ecological stoichiometry. In the ocean, when averaged over a large number of species, an overall convergence of stoichiometry seems to exist in phytoplankton approaching the C:N:P values of 106:16:1 proposed by Redfield, coincident with that of dissolved nutrients in deep waters (Redfield 1960; see also Falkowski 2000; Klausmeier *et al*., 2004, 2008). The macronutrients C, N, P, as well as S are the major constituents of organic macromolecules of cells (i.e., proteins, nucleic acids, carbohydrates and lipids) on which fundamental structures and functions depend (e.g. Ríos & Fraga, 1987; Giordano, 2013). Changes in macronutrient quotas modify the organic composition of organisms and hence their stoichiometry, e.g., when N, P and S quotas decrease, C is reallocated to organic pools that do not contain these elements, such as carbohydrates and lipids (Heldal *et al*., 1996; Geider & La Roche, 2002; Klausmeier 2004, 2008). The reallocation of C is governed by genotypic (Palmucci *et al*., 2011), energetic (Norici *et al*., 2011; Fanesi *et al*., 2014; Ruan & Giordano, 2017), spatial (Palmucci *et al*., 2011), as well as structural and functional constraints (Giordano, 2001; Montechiaro *et al*., 2006; Kaffes *et al*., 2010). In addition, these changes can be buffered if the cells are able to store specific macroelements (e.g. P as polyphosphate) or uptake those elements more than required (i.e. luxury uptake, John & Flynn, 2000; Docampo *et al*., 2005; Eide, 2006; Marchetti *et al*., 2009; Nuester *et al*., 2012; Roh *et al*., 2012).

So, even though there is variability at the species level, when they are aggregated into higher taxonomic groupings evidence of taxa-specific elemental fingerprints emerge (Ho *et al*., 2003; Quigg *et al.,*2003, 2011; Segura-Noguera *et al*., 2016). It has been proposed that these fingerprints reflect differences in the origin of the plastids acquired by eukaryotic algae (plastid imprint hypothesis, Quigg *et al*., 2011), giving rise to the “red” and “green” lineages (Grzebyk *et al*., 2003; Bodył, 2017). In addition, changes in environmental conditions and in the physiological status of cells like temperature (Moorthi *et al*., 2016), irradiance (Finkel *et al*., 2006; Thrane *et al*., 2016), nutrient availability (Rhee & Gotham, 1980; Bi *et al*., 2012; Palmucci & Giordano, 2012; Segura-Noguera *et al*., 2016; Ruan & Giordano, 2017), growth rate (Droop, 1974; Goldman *et al*., 1979; Burkhardt *et al.,*1999; Giordano, 2013; Hong-Hermesdorf *et al*., 2014) and size (Finkel *et al*., 2010; Garcia *et al*., 2016, Segura-Noguera *et al*., 2016) also influence cellular elemental composition. There must be, however, inherent constraints to this stoichiometric flexibility, since basic structural and metabolic requirements, that are shared across taxa, must be satisfied (Rhee, 1978; Geider & La Roche, 2002; Loladze & Elser, 2011).

Micronutrient cellular quotas display significantly greater variability compared to macronutrients (Giordano, 2013). Nevertheless, distinct patterns in the relative abundance of micronutrients emerge within cultured species. Ho *et al*. (2003), for instance, suggested an element abundance hierarchy in cultured marine phytoplankton species: Fe>Mn>Zn>Cu>Co≈Cd>Mo, while Twining & Baines (2013) presented a comparable hierarchy for natural phytoplankton assemblages, differing only in the positions of Zn and Mn relative to Ho *et al*. (2003): Fe≈Zn>Mn≈Ni≈Cu>>Co≈Cd. A variety of cellular mechanisms can influence the stoichiometrical relationships of micronutrients, for example: 1) some metallo-proteins can associate to alternate ligands under certain conditions (Price & Morel, 1990); 2) entirely different proteins can fill in a specific function, e.g. if the primary metallo-protein does not have its ligand available (La Roche *et al*., 1996); and 3) different elements may compete for the same transport system (Sunda & Huntsman, 1998). Quigg *et al*. (2003) suggested that algae of the red and green lineages are inherently different with respect to their metal utilization and content, with the red lineage being prone to favor metals that are more abundant in fully oxygenated waters.

Sulfur is a macronutrient often excluded from stoichiometric studies possibly since this element is rarely limiting in modern oceans. Yet, with a cell quota similar and sometimes higher than that of P (e.g. Giordano, 2013; Segura-Noguera *et al*., 2016), S has crucial functions in the cell: 1) S is contained in cysteine, glutathione and phytochelatins forming part of thiol groups, which are involved in protein tertiary structures, redox enzyme regulation, and detoxification of heavy metals and oxidants; 2) S is also found in abundant multifunctional compounds like DMSP, and in sulfolipids (Giordano & Raven, 2014; Giordano & Prioretti, 2016); and 3) recent studies have shown that S availability influences the rate of peptide elongation on ribosomes (Giordano *et al*., 2015). Similarly to other macronutrients, changes in S abundance and stoichiometry and its consequent variations in the organic composition of algal cells has unavoidable reverberation in the trophic web, both in extant aquatic environments and in long-term ecological trajectories. Beyond the plastid imprint hypothesis (Quigg *et al*., 2011), which suggests that red lineage species dominate due to their plastid content, it is proposed that Paleozoic-Mesozoic chemical shifts, particularly the increased environmental S concentration, favored Mesozoic red lineage algae with higher S requirements and heightened metabolic S fluxes (Giordano *et al*., 2005; Norici *et al*., 2005; Ratti *et al*., 2011, 2013; Prioretti & Giordano, 2016). Furthermore, it has been hypothesized that the rise in the environmental S concentration amplified the aggressiveness of grazers, which elicited clade-specific responses and ultimately shaped the phytoplankton clade composition of modern ocean (Giordano *et al*., 2018).

Limited research has addressed the effect of S availability on cell stoichiometry. These studies, encompassing a restricted range of elements and representative species, show that sulfate availability can affect both macro- and microelement cell quotas. For instance, the diatom *Thalassiosira weissflogii* cultured at 5 mM sulfate had lower C:P, N:P and S:P ratio than their 30 mM grown counterparts, however, the same did not occur in the cyanobacterium *Synechococcus* sp. or in the green alga *Tetraselmis suecica* (Prioretti & Giordano 2016, Ratti *et al*., 2013). Sulfate concentration in the media also influenced the cell quotas of metals such as Fe, Mn, and Cu in the diatom *Thalassiosira pseudonana*, while no major impact of S availability was observed on the elemental composition of *Synechococcus* sp, *Tetraselmis suecica* and the dinoflagellate *Amphidinium klebsii* (Prioretti & Giordano, 2016). Unfortunately, these few studies do not allow us to unravel the potential divergent response in the elemental composition among major taxonomic groups derived from sulfate availability.

In the ocean and other bodies of water, chemical changes occur in response to acute and short/medium-term perturbations (such as pollution), but also as a result of fluctuations that occurred over the history of the planet (Giordano *et al*., 2005, and references therein; Morel *et al*., 2014). In this study, we focus on the elemental stoichiometry of phytoplankton under different sulfate availabilities in the ocean and explore how these observed changes might correlate with cell metabolism and potentially relate to evolutionary trajectories of species within the red and green lineages. To do so, we studied the influence of S availability on the elemental concentration and cell stoichiometry of 20 phytoplankton species, including 11 from the red lineage, 7 from the green lineage, and 2 cyanobacteria, and we examine patterns in the stoichiometry of major and trace elements as a first step to identifying common cellular mechanisms relating to compensation for changes in S availability. To simulate conditions, we employed 5 mM sulfate to mirror early oceans and 28 mM sulfate to mimic modern oceanic environments (Ratti *et al*., 2011; Giordano & Raven, 2014). Using the same medium, AMCONA, facilitated consistent growth across a broad range of phytoplankton species, mitigating potential artifacts arising from different growth media.

## Materials and Methods

### Culture conditions and determination of specific growth rate and cell volume

All species in this study (Table 1) were received from the Culture Collection of Algae and Protozoa (CCAP) and were grown in Artificial Multipurpose COmplement for the Nutrition of Algae (AMCONA) medium (Fanesi *et al*., 2014), in wide-necked 250 ml erlenmeyer flasks containing 200 ml of culture. The sulfate (provided as Na_2_SO_4_) concentration in the medium was modified from the original recipe to obtain 28 mM (corresponding to that of today’s ocean, Norici *et al*., 2005) and 5 mM (approximately corresponding to the early Paleozoic concentration, Ratti *et al*., 2011, 2013; Giordano *et al*., 2018). The total osmolarity of the medium was adjusted to the same value with NaCl. All cultures were maintained in a growth chamber (Perani FT 1500), at a constant temperature of 20 °C, under continuous light and a photon flux density (PFD) of 110 µmol photons m^-2^s^-1^ provided by cool white fluorescent tubes. For each species, the specific growth rate was determined from the exponential portion of the growth curve of batch cultures, according to Monod’s equation. Semi-continuous cultures were established with daily dilution rates imposed to match the maximum growth rate. All semi-continuous cultures were grown for at least six generations prior to sampling. For each species and S condition, three replicate flasks were used, and three samplings were taken from each flask. Each sampling was separated in time by at least one generation. Cell counts were verified daily with an automatic cell counter (CASY TT, Innovatis AG, Reutlingen, Germany).

**Table 1.** List of the species studied, including the lineage, taxonomic classification, cellular volume (μm^3^), dry weight (pg) and growth rates (day^-1^) of monocultures grown with different S concentrations (5 or 28 mM sulfate) in the media.

| Lineage | Classification |  | Strain | 5 mM sulfate |  |  | 28 mM sulfate |  |  |
| --- | --- | --- | --- | --- | --- | --- | --- | --- | --- |
| | Phylum | Class | | V ($\mu\text{m}^3$) | dry weight (pg) | growth rate | V ($\mu\text{m}^3$) | dry weight (pg) | growth rate |
| Green lineage<br>Chl *a* + *b* | Chlorophyta | Chlorophyceae | *Asteromonas gracialis* | 2097.6 ± 142.3 | 655.4 ± 83.5 | 0.188 | 2043.7 ± 139.7 | 642.5 ± 140.2 | 0.235 |
| | Chlorophyta | Chlorophyceae | *Coccomyxa parasitica* | 37.4 ± 1.9 | 14.4 ± 2.3 | 0.593 | 39.3 ± 3.5 | 15.3 ± 3.3 | 0.518 |
| | Chlorophyta | Chlorophyceae | *Dunaliella bardawil* | 1038.6 ± 71.0 | 284.9 ± 24.3 | 0.414 | 1049.9 ± 47.1 | 301.9 ± 50.0 | 0.607 |
| | Chlorophyta | Chlorophyceae | *Dunaliella salina* | 153.6 ± 13.9 | 41.7 ± 4.3 | 1.003 | 153.6 ± 9.0 | 33.1 ± 4.1 | 1.040 |
| | Chlorophyta | Chlorophyceae | *Dunaliella tertiolecta* | 227.2 ± 11.9 | 66.8 ± 6.9 | 1.301 | 229.8 ± 6.8 | 65.1 ± 5.8 | 1.328 |
| | Chlorophyta | Trebouxiophyceae | *Nannochloris atomus* | 40.0 ± 1.5 | 12.9 ± 1.8 | 0.693 | 40.1 ± 0.7 | 14.6 ± 1.1 | 0.699 |
| | Chlorophyta | Chlorodendrophyceae | *Tetraselmis suecica* | 406.1 ± 9.0 | 84.0 ± 8.5 | 0.737 | 415.2 ± 7.6 | 96.6 ± 7.5 | 1.003 |
| Red lineage<br>Chl *a* + *c* | Haptophyta | Pavlovophyceae | *Diacronema lutheri* | 45.4 ± 2.6 | 7.9 ± 1.8 | 0.504 | 47.1 ± 3.9 | 9.7 ± 1.4 | 0.494 |
| | Ocrophyta | Eustigmatophyceae | *Nannochloropsis gaditana* | 19.3 ± 2.1 | 3.2 ± 0.7 | 0.921 | 18.7 ± 2.1 | 3.4 ± 0.7 | 0.856 |
| | Ocrophyta | Eustigmatophyceae | *Nannochloropsis salina* | 21.6 ± 1.1 | 4.3 ± 1.2 | 0.760 | 22.4 ± 16 | 3.7 ± 0.5 | 0.639 |
| | Ocrophyta | Bacillariophyceae | *Fragilaria pinnata* | 241.8 ± 27.6 | 84.2 ± 9.1 | 0.750 | 253.2 ± 27.7 | 99.4 ± 11.5 | 0.767 |
| | Ocrophyta | Bacillariophyceae | *Phaeodactylum tricornutum* | 59.0 ± 4.2 | 9.5 ± 1.7 | 0.505 | 58.3 ± 3.9 | 9.8 ± 1.9 | 0.783 |
| | Ocrophyta | Bacillariophyceae | *Thalassiosira pseudonana* | 34.5 ± 2.2 | 8.0 ± 0.9 | 0.819 | 36.4 ± 1.9 | 8.8 ± 1.8 | 0.994 |
| | Ocrophyta | Bacillariophyceae | *Thalassiosira weissflogii* | 469.9 ± 20.9 | 93.6 ± 13.9 | 1.179 | 463.9 ± 21.4 | 95.8 ± 15.8 | 1.108 |
| | Rhodophyta | Porphyridiophyceae | *Porphyridium purpureum* | 159.2 ± 16.8 | 60.4 ± 13.0 | 1.326 | 178.4 ± 9.5 | 75.7 ± 15.2 | 1.258 |
| | Cryptophyta | Cryptophyceae | *Rhinomonas reticulata* var. *reticulata* | 597.8 ± 33.3 | 72.5 ± 9.6 | 0.468 | 627.6 ± 42.9 | 70.9 ± 8.4 | 0.472 |
| | Myzozoa | Dinophyceae | *Amphidinium carterae* | 878.4 ± 31.2 | 105.8 ± 20.2 | 0.494 | 971.0 ± 22.7 | 118.2 ± 11.9 | 0.532 |
| | Myzozoa | Dinophyceae | *Amphidinium klebsii* | 1060.3 ± 53.2 | 157.6 ± 20.4 | 0.531 | 1054.5 ± 53.3 | 160.1 ± 16.4 | 0.765 |
| Cyanobacteria<br>Chl *a* +<br>phycocyanin | Cyanobacteria | Cyanophyceae | *Arthrospira brevis* | 4.5 ± 0.1 | 1.5 ± 0.1 | 0.438 | 6.1 ± 0.3 | 1.8 ± 0.2 | 0.602 |
| | Cyanobacteria | Cyanophyceae | *Synechococcus* sp. | 8.5 ± 0.3 | 2.9 ± 0.7 | 1.007 | 8.1 ± 0.2 | 2.8 ± 0.6 | 0.775 |

### Elemental analysis

The C and N cell quotas were determined with an elemental combustion system ECS CHNSO 4010 (Costech Analytical Technologies Inc., Milan, Italy). Samples were prepared by washing cells twice with an iso-osmotic ammonium formate solution relative to the culture media; the cell pellet was then dried at 100°C until it reached a stablized weight. Biomass aliquots ranging from 0.5 to 5 mg of dry weight were used for analysis, and, in each, a small amount of vanadium peroxide was added to facilitate combustion. Sulfanilamide (C:N:S = 6:2:1) was used as standard. The oven temperature was set to 980°C, the chromatographic column temperature was set to 70°C and the air pressure was set to three bar. Gas fluxes were set to 100 ml min^-1^ for Helium, and 30 ml min^-1^ for oxygen. Data acquisition and subsequent analysis were performed with the software EAS-CLARITY (DataApex Ltd. 2006, Prague, Czech Republic).

The elemental cell quotas of Si, P, S, K, Ca, Mn, Fe, Zn, Cr, Co, Cu, Se, Br and Sr were measured by a Picofox S2 total reflection X-ray fluorescence (TXRF) spectrometer (Bruker AXS Microanalysis GmbH, Ettlingen, Germany). To ensure precise analysis, a sample preparation protocol was conducted set up to remove non-cellular elemental traces. All materials for sample preparation and elemental analysis underwent acid-washing in 10% trace-metal grade hydrochloric acid at 60°C for at least 12 hours, followed by thoroughly rinsing in Milli-Q water. Considering the propensity for ambient Fe to adsorb onto the cell surface (Morel & Hering, 1993), we designed a procedure to wash the cells prior to analysis. The washing reagent Oxalate-EDTA (OE) was prepared based on the method proposed by Tovar-Sanchez *et al*. (2003): for a final volume of 100 ml, 1.95 grams EDTA-Na_2_·2H_2_O and 0.5 grams NaCl were first added to about 60 ml milli-Q water. This solution was then brought to pH 6.5 with 10M NaOH. After, 1.26 grams of oxalic acid (C_2_H_2_O_4_·2H_2_O) was dissolved in the solution, and the pH was adjusted to 6.5 with 10M NaOH. The resulting solution was brought to 100 ml volume and stored at 4°C for a maximum of 60 days.

The samples were harvested and washed as follows: first, the cells in the semi-continuous cultures were counted; a suitable volume of the culture (usually 10 ml) was then centrifuged at between 2000 and 4000 rpm depending on the size and fragility of the species. To avoid losing cells, 1 ml of the supernatant was left on top of the pellet, and 3 ml of the OE solution at room temperature (20°C) was then added. After resuspension at room temperature for 5 minutes, the cells were centrifuged and suspended in OE again. A third wash with a trace-metal-clean 0.5M NaCl solution was performed. Finally, all supernatant was carefully removed and the pellet was re-suspended in 995 µl (990µl for large pellets) milli-Q water. Five microliters of gallium standard solution were then added to bring the final sample volume to exactly 1000 µl. Care was taken at this point to thoroughly and homogeneously re-suspend the pellet with vigorous vortexing; when necessary a sonicating bath and freeze-thaw cycles were used. Ten microliters of the samples were pipetted onto quartz sample holder discs. Acrylic sample holder discs were used for diatoms to avoid the interference of quartz with the measurement of silica. In the case of quartz discs, the sample was dried at 60°C, while temperature sensitive acrylic discs were dried at room temperature, which took about 20 minutes. Samples were then analyzed in a Picofox S2 spectrometer (Bruker AXS Microanalysis GmbH, Berlin, Germany). Samples readings were conducted for 1000 seconds. Data acquisition was performed with the software SPECTRA 5.3 (Bruker AXS Microanalysis GmbH).

### Data Analysis

Three biological replicates were used for samples analyzed by gas chromatography in the elemental combustion system, while nine biological replicates were employed for analysis by TXRF spectrometry. All elemental data were converted to cellular quotas and molar concentrations. Statistical analyses were performed using R Software (v.4.2.2, R Core Team, 2022). Differences between mean values were compared after testing the normality of the distributions (the Shapiro-Wilk normality test) and the homogeneity of variances (the Levene’s Test of Equality of Variances in lawstat package, Gastwirth *et al*., 2023). Variables not normally distributed were log-transformed. Student’s two-tailed t-tests for paired samples with 1, 6, and 10 degrees of freedom for cyanobaceria, green lineage and red lineage, respectively, were used to discover differences within lineages grown with different media sulfate concentrations. To reveal differences between lineages, Student’s two-tailed t-tests were applied. Si was only detected in diatoms, and it was not detectable in one of the species grown at 28 mM, so the statistical tests for this element have two degrees of freedom.

## Results

### Effect of S availability on growth rate, cell volume and dry weight

We studied seven species of microalgae belonging to the green lineage, eleven to the red lineage, and two cyanobacteria. The taxonomic group, dry weight and volume per cell, as well as growth rate are listed in Table 1. Within each selection of lineages, we had a wide range of cell sizes and growth rates. The species with the highest growth rate was *Porphyridium purpurea* (belonging to the red lineage), followed closely by *Dunaliella tertiolecta* (green lineage). The species with relatively slower growth rates often corresponded to those with larger cell volumes (e.g., *Asteromonas graciales* and *Rhinomonas reticulata*), however this relationship did not always hold. The two cyanobacteria, for instance, showed a large difference in growth rate at 5 mM sulfate despite being very similar in size. Most species had very similar volumes and maximum growth rates in media with low and high S concentrations, and the largest rates found in six out of seven species of the green lineage, and in six out of 11 species of the red lineage were not statistically significant (*p* > 0.05) (Table 1). On average, the dry weight of cells from the red lineage significantly increased when grown under high sulfate conditions (6.4 ± 10.3%, *p* ≤ 0.05), but not for the green lineage (Table 1). We did not detect statistically significant differences in volume, dry weight or growth rate between cells from the green and the red lineage grown with the same sulfate concentration. The volume of cyanobacteria was lower than that of the red and green lineages (*p* ≤ 0.05), as was as dry weight (*p* ≤ 0.06 and *p* ≤ 0.01 for green and red lineage, respectively), for both sulfate concentrations.

### Elemental composition and effects of S availability on elemental cell quotas and concentrations

The variability of C, N, P and S molar concentrations, as a function of the species and sulfate concentration in the media, was 35% for C, 40% for N, 45% for P, and 70% for S (Table 2). The variability for most of the remaining elements was between 40-89%, with Sr being the most variable (coefficient of variation (CV) = 168% at 5 mM sulfate, and 135% at 28 mM sulfate, or 95% when *D. salina* is not included). C and N concentrations were higher in the green lineage and cyanobacteria compared to the red lineage (*p* ≤ 0.01) in both sulfate conditions. Cyanobacteria had the highest P concentration (*p* ≤ 0.05), while the P concentration was similar between the green and red lineage (*p* > 0.2). The red lineage algae had almost double the S cell concentration (*p* ≤ 0.05) compared to the green lineage algae (Table 2, Figure 1). Si was only detected in diatoms, and, when *P. tricornutum* was removed from the analysis due to its unusual low Si quota (Lewin *et al*., 1958), its concentration was similar between *F. pinnata and T. pseudonana* (Table 2). Selenium was detected only in the two dinoflagellates with a variability of 40% (Table 2). None of the species studied contained detectable levels of Co.

**Figure 1.**
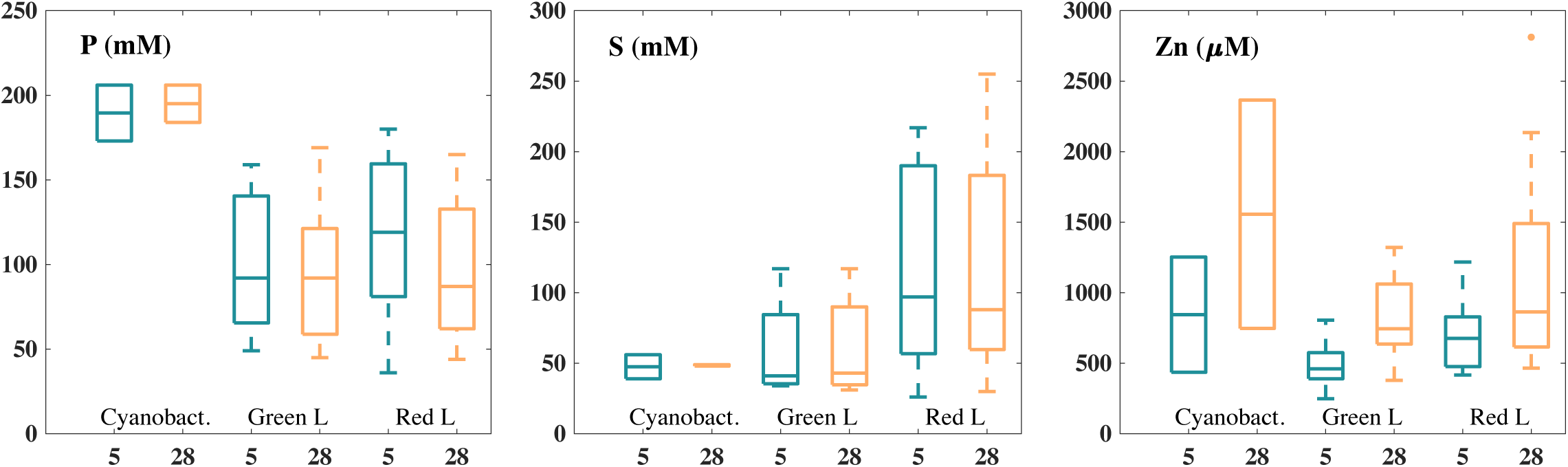
Boxplots of P (mM), S (mM) and Zn (µM) concentration in Cyanobacteria (n = 2), in species from the Green (n = 7) and from the Red (n = 11) Lineages grown in 5 mM (orange) or 28 mM (blue) of sulfate concentration in the media. Note that y-axes change.

**Table 2.**
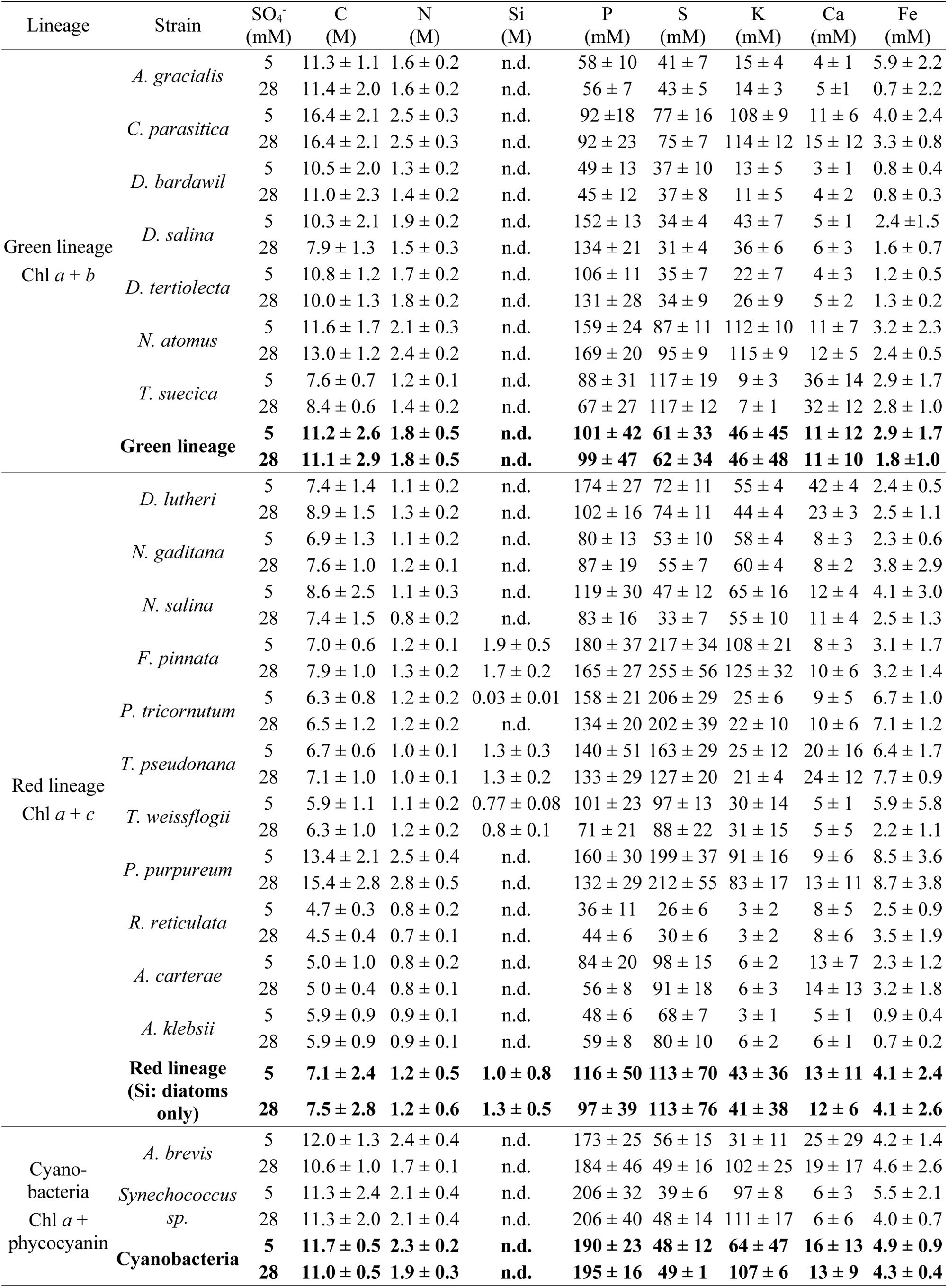
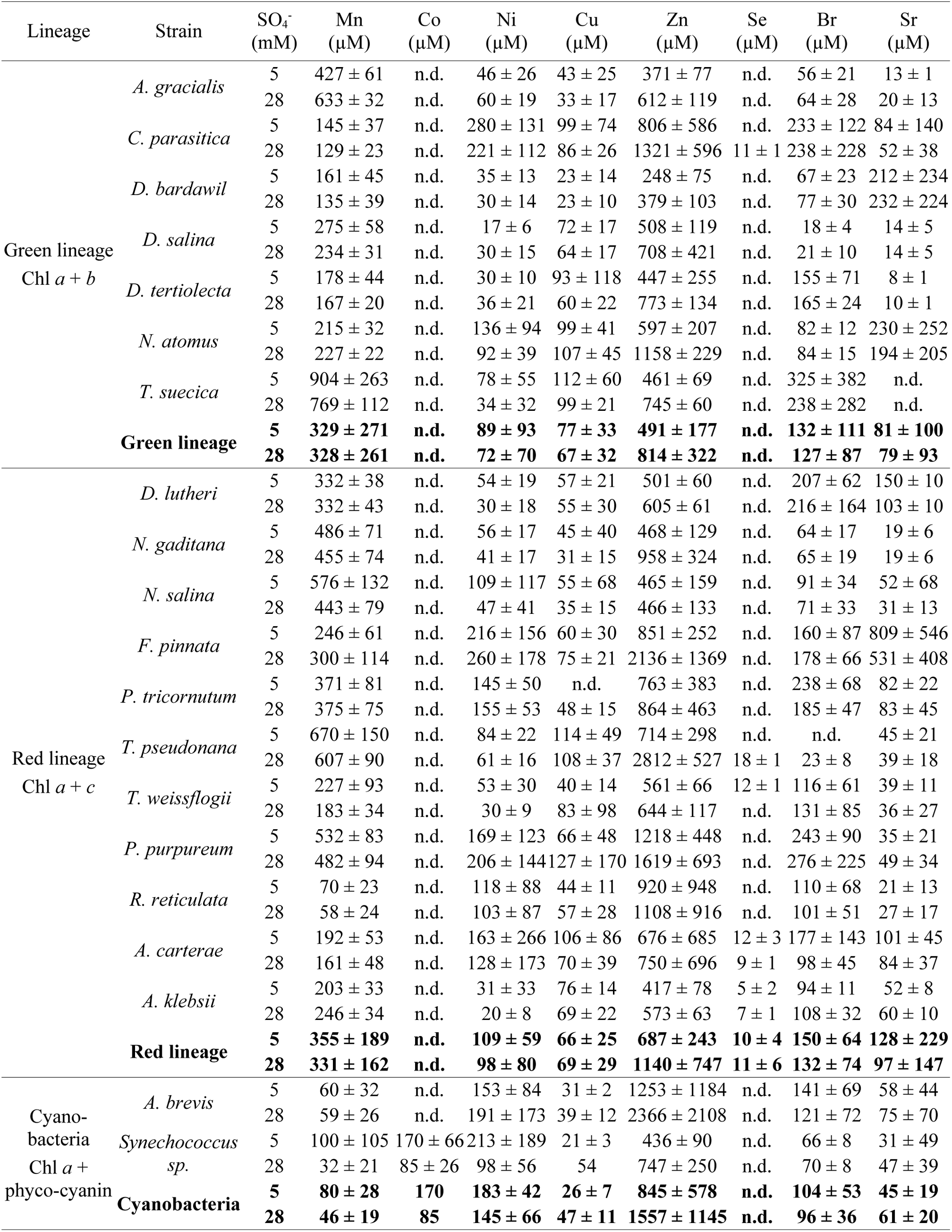
Molar concentration (average ± standard deviation) in the species studied grown at 5 or 28 mM of sulfate in the media. Note that the units of the concentrations vary between elements. n.d. = not detected. Bold numbers: average of the lineage or of the cyanobacteria.

| Lineage | Strain | SO<sub>4</sub><sup>-</sup><br>(mM) | C<br>(M) | N<br>(M) | Si<br>(M) | P<br>(mM) | S<br>(mM) | K<br>(mM) | Ca<br>(mM) | Fe<br>(mM) |
| --- | --- | --- | --- | --- | --- | --- | --- | --- | --- | --- |
| Green lineage<br>Chl *a* + *b* | *A. gracialis* | 5 | 11.3 ± 1.1 | 1.6 ± 0.2 | n.d. | 58 ± 10 | 41 ± 7 | 15 ± 4 | 4 ± 1 | 5.9 ± 2.2 |
|  |  | 28 | 11.4 ± 2.0 | 1.6 ± 0.2 | n.d. | 56 ± 7 | 43 ± 5 | 14 ± 3 | 5 ± 1 | 0.7 ± 2.2 |
|  | *C. parasitica* | 5 | 16.4 ± 2.1 | 2.5 ± 0.3 | n.d. | 92 ± 18 | 77 ± 16 | 108 ± 9 | 11 ± 6 | 4.0 ± 2.4 |
|  |  | 28 | 16.4 ± 2.1 | 2.5 ± 0.3 | n.d. | 92 ± 23 | 75 ± 7 | 114 ± 12 | 15 ± 12 | 3.3 ± 0.8 |
|  | *D. bardawil* | 5 | 10.5 ± 2.0 | 1.3 ± 0.2 | n.d. | 49 ± 13 | 37 ± 10 | 13 ± 5 | 3 ± 1 | 0.8 ± 0.4 |
|  |  | 28 | 11.0 ± 2.3 | 1.4 ± 0.2 | n.d. | 45 ± 12 | 37 ± 8 | 11 ± 5 | 4 ± 2 | 0.8 ± 0.3 |
|  | *D. salina* | 5 | 10.3 ± 2.1 | 1.9 ± 0.2 | n.d. | 152 ± 13 | 34 ± 4 | 43 ± 7 | 5 ± 1 | 2.4 ± 1.5 |
|  |  | 28 | 7.9 ± 1.3 | 1.5 ± 0.3 | n.d. | 134 ± 21 | 31 ± 4 | 36 ± 6 | 6 ± 3 | 1.6 ± 0.7 |
|  | *D. tertiolecta* | 5 | 10.8 ± 1.2 | 1.7 ± 0.2 | n.d. | 106 ± 11 | 35 ± 7 | 22 ± 7 | 4 ± 3 | 1.2 ± 0.5 |
|  |  | 28 | 10.0 ± 1.3 | 1.8 ± 0.2 | n.d. | 131 ± 28 | 34 ± 9 | 26 ± 9 | 5 ± 2 | 1.3 ± 0.2 |
|  | *N. atomus* | 5 | 11.6 ± 1.7 | 2.1 ± 0.3 | n.d. | 159 ± 24 | 87 ± 11 | 112 ± 10 | 11 ± 7 | 3.2 ± 2.3 |
|  |  | 28 | 13.0 ± 1.2 | 2.4 ± 0.2 | n.d. | 169 ± 20 | 95 ± 9 | 115 ± 9 | 12 ± 5 | 2.4 ± 0.5 |
|  | *T. suecica* | 5 | 7.6 ± 0.7 | 1.2 ± 0.1 | n.d. | 88 ± 31 | 117 ± 19 | 9 ± 3 | 36 ± 14 | 2.9 ± 1.7 |
|  |  | 28 | 8.4 ± 0.6 | 1.4 ± 0.2 | n.d. | 67 ± 27 | 117 ± 12 | 7 ± 1 | 32 ± 12 | 2.8 ± 1.0 |
|  | **Green lineage** | **5** | **11.2 ± 2.6** | **1.8 ± 0.5** | **n.d.** | **101 ± 42** | **61 ± 33** | **46 ± 45** | **11 ± 12** | **2.9 ± 1.7** |
|  |  | **28** | **11.1 ± 2.9** | **1.8 ± 0.5** | **n.d.** | **99 ± 47** | **62 ± 34** | **46 ± 48** | **11 ± 10** | **1.8 ± 1.0** |
| Red lineage<br>Chl *a* + *c* | *D. lutheri* | 5 | 7.4 ± 1.4 | 1.1 ± 0.2 | n.d. | 174 ± 27 | 72 ± 11 | 55 ± 4 | 42 ± 4 | 2.4 ± 0.5 |
|  |  | 28 | 8.9 ± 1.5 | 1.3 ± 0.2 | n.d. | 102 ± 16 | 74 ± 11 | 44 ± 4 | 23 ± 3 | 2.5 ± 1.1 |
|  | *N. gaditana* | 5 | 6.9 ± 1.3 | 1.1 ± 0.2 | n.d. | 80 ± 13 | 53 ± 10 | 58 ± 4 | 8 ± 3 | 2.3 ± 0.6 |
|  |  | 28 | 7.6 ± 1.0 | 1.2 ± 0.1 | n.d. | 87 ± 19 | 55 ± 7 | 60 ± 4 | 8 ± 2 | 3.8 ± 2.9 |
|  | *N. salina* | 5 | 8.6 ± 2.5 | 1.1 ± 0.3 | n.d. | 119 ± 30 | 47 ± 12 | 65 ± 16 | 12 ± 4 | 4.1 ± 3.0 |
|  |  | 28 | 7.4 ± 1.5 | 0.8 ± 0.2 | n.d. | 83 ± 16 | 33 ± 7 | 55 ± 10 | 11 ± 4 | 2.5 ± 1.3 |
|  | *F. pinnata* | 5 | 7.0 ± 0.6 | 1.2 ± 0.1 | 1.9 ± 0.5 | 180 ± 37 | 217 ± 34 | 108 ± 21 | 8 ± 3 | 3.1 ± 1.7 |
|  |  | 28 | 7.9 ± 1.0 | 1.3 ± 0.2 | 1.7 ± 0.2 | 165 ± 27 | 255 ± 56 | 125 ± 32 | 10 ± 6 | 3.2 ± 1.4 |
|  | *P. tricornutum* | 5 | 6.3 ± 0.8 | 1.2 ± 0.2 | 0.03 ± 0.01 | 158 ± 21 | 206 ± 29 | 25 ± 6 | 9 ± 5 | 6.7 ± 1.0 |
|  |  | 28 | 6.5 ± 1.2 | 1.2 ± 0.2 | n.d. | 134 ± 20 | 202 ± 39 | 22 ± 10 | 10 ± 6 | 7.1 ± 1.2 |
|  | *T. pseudonana* | 5 | 6.7 ± 0.6 | 1.0 ± 0.1 | 1.3 ± 0.3 | 140 ± 51 | 163 ± 29 | 25 ± 12 | 20 ± 16 | 6.4 ± 1.7 |
|  |  | 28 | 7.1 ± 1.0 | 1.0 ± 0.1 | 1.3 ± 0.2 | 133 ± 29 | 127 ± 20 | 21 ± 4 | 24 ± 12 | 7.7 ± 0.9 |
|  | *T. weissflogii* | 5 | 5.9 ± 1.1 | 1.1 ± 0.2 | 0.77 ± 0.08 | 101 ± 23 | 97 ± 13 | 30 ± 14 | 5 ± 1 | 5.9 ± 5.8 |
|  |  | 28 | 6.3 ± 1.0 | 1.2 ± 0.2 | 0.8 ± 0.1 | 71 ± 21 | 88 ± 22 | 31 ± 15 | 5 ± 5 | 2.2 ± 1.1 |
|  | *P. purpureum* | 5 | 13.4 ± 2.1 | 2.5 ± 0.4 | n.d. | 160 ± 30 | 199 ± 37 | 91 ± 16 | 9 ± 6 | 8.5 ± 3.6 |
|  |  | 28 | 15.4 ± 2.8 | 2.8 ± 0.5 | n.d. | 132 ± 29 | 212 ± 55 | 83 ± 17 | 13 ± 11 | 8.7 ± 3.8 |
|  | *R. reticulata* | 5 | 4.7 ± 0.3 | 0.8 ± 0.2 | n.d. | 36 ± 11 | 26 ± 6 | 3 ± 2 | 8 ± 5 | 2.5 ± 0.9 |
|  |  | 28 | 4.5 ± 0.4 | 0.7 ± 0.1 | n.d. | 44 ± 6 | 30 ± 6 | 3 ± 2 | 8 ± 6 | 3.5 ± 1.9 |
|  | *A. carterae* | 5 | 5.0 ± 1.0 | 0.8 ± 0.2 | n.d. | 84 ± 20 | 98 ± 15 | 6 ± 2 | 13 ± 7 | 2.3 ± 1.2 |
|  |  | 28 | 5.0 ± 0.4 | 0.8 ± 0.1 | n.d. | 56 ± 8 | 91 ± 18 | 6 ± 3 | 14 ± 13 | 3.2 ± 1.8 |
|  | *A. klebsii* | 5 | 5.9 ± 0.9 | 0.9 ± 0.1 | n.d. | 48 ± 6 | 68 ± 7 | 3 ± 1 | 5 ± 1 | 0.9 ± 0.4 |
|  |  | 28 | 5.9 ± 0.9 | 0.9 ± 0.1 | n.d. | 59 ± 8 | 80 ± 10 | 6 ± 2 | 6 ± 1 | 0.7 ± 0.2 |
|  | **Red lineage**<br>**(Si: diatoms**<br>**only)** | **5** | **7.1 ± 2.4** | **1.2 ± 0.5** | **1.0 ± 0.8** | **116 ± 50** | **113 ± 70** | **43 ± 36** | **13 ± 11** | **4.1 ± 2.4** |
|  |  | **28** | **7.5 ± 2.8** | **1.2 ± 0.6** | **1.3 ± 0.5** | **97 ± 39** | **113 ± 76** | **41 ± 38** | **12 ± 6** | **4.1 ± 2.6** |
| Cyano-<br>bacteria<br>Chl *a* +<br>phycocyanin | *A. brevis* | 5 | 12.0 ± 1.3 | 2.4 ± 0.4 | n.d. | 173 ± 25 | 56 ± 15 | 31 ± 11 | 25 ± 29 | 4.2 ± 1.4 |
|  |  | 28 | 10.6 ± 1.0 | 1.7 ± 0.1 | n.d. | 184 ± 46 | 49 ± 16 | 102 ± 25 | 19 ± 17 | 4.6 ± 2.6 |
|  | *Synechococcus*<br>*sp.* | 5 | 11.3 ± 2.4 | 2.1 ± 0.4 | n.d. | 206 ± 32 | 39 ± 6 | 97 ± 8 | 6 ± 3 | 5.5 ± 2.1 |
|  |  | 28 | 11.3 ± 2.0 | 2.1 ± 0.4 | n.d. | 206 ± 40 | 48 ± 14 | 111 ± 17 | 6 ± 6 | 4.0 ± 0.7 |
|  | **Cyanobacteria** | **5** | **11.7 ± 0.5** | **2.3 ± 0.2** | **n.d.** | **190 ± 23** | **48 ± 12** | **64 ± 47** | **16 ± 13** | **4.9 ± 0.9** |
|  |  | **28** | **11.0 ± 0.5** | **1.9 ± 0.3** | **n.d.** | **195 ± 16** | **49 ± 1** | **107 ± 6** | **13 ± 9** | **4.3 ± 0.4** |

**Table 2. (cont.)**
| Lineage | Strain | SO<sub>4</sub><sup>-</sup><br>(mM) | Mn<br>(μM) | Co<br>(μM) | Ni<br>(μM) | Cu<br>(μM) | Zn<br>(μM) | Se<br>(μM) | Br<br>(μM) | Sr<br>(μM) |
| --- | --- | --- | --- | --- | --- | --- | --- | --- | --- | --- |
| Green lineage<br>Chl *a* + *b* | *A. gracialis* | 5 | 427 ± 61 | n.d. | 46 ± 26 | 43 ± 25 | 371 ± 77 | n.d. | 56 ± 21 | 13 ± 1 |
|  |  | 28 | 633 ± 32 | n.d. | 60 ± 19 | 33 ± 17 | 612 ± 119 | n.d. | 64 ± 28 | 20 ± 13 |
|  | *C. parasitica* | 5 | 145 ± 37 | n.d. | 280 ± 131 | 99 ± 74 | 806 ± 586 | n.d. | 233 ± 122 | 84 ± 140 |
|  |  | 28 | 129 ± 23 | n.d. | 221 ± 112 | 86 ± 26 | 1321 ± 596 | 11 ± 1 | 238 ± 228 | 52 ± 38 |
|  | *D. bardawil* | 5 | 161 ± 45 | n.d. | 35 ± 13 | 23 ± 14 | 248 ± 75 | n.d. | 67 ± 23 | 212 ± 234 |
|  |  | 28 | 135 ± 39 | n.d. | 30 ± 14 | 23 ± 10 | 379 ± 103 | n.d. | 77 ± 30 | 232 ± 224 |
|  | *D. salina* | 5 | 275 ± 58 | n.d. | 17 ± 6 | 72 ± 17 | 508 ± 119 | n.d. | 18 ± 4 | 14 ± 5 |
|  |  | 28 | 234 ± 31 | n.d. | 30 ± 15 | 64 ± 17 | 708 ± 421 | n.d. | 21 ± 10 | 14 ± 5 |
|  | *D. tertiolecta* | 5 | 178 ± 44 | n.d. | 30 ± 10 | 93 ± 118 | 447 ± 255 | n.d. | 155 ± 71 | 8 ± 1 |
|  |  | 28 | 167 ± 20 | n.d. | 36 ± 21 | 60 ± 22 | 773 ± 134 | n.d. | 165 ± 24 | 10 ± 1 |
|  | *N. atomus* | 5 | 215 ± 32 | n.d. | 136 ± 94 | 99 ± 41 | 597 ± 207 | n.d. | 82 ± 12 | 230 ± 252 |
|  |  | 28 | 227 ± 22 | n.d. | 92 ± 39 | 107 ± 45 | 1158 ± 229 | n.d. | 84 ± 15 | 194 ± 205 |
|  | *T. suecica* | 5 | 904 ± 263 | n.d. | 78 ± 55 | 112 ± 60 | 461 ± 69 | n.d. | 325 ± 382 | n.d. |
|  |  | 28 | 769 ± 112 | n.d. | 34 ± 32 | 99 ± 21 | 745 ± 60 | n.d. | 238 ± 282 | n.d. |
|  | **Green lineage** | **5** | **329 ± 271** | **n.d.** | **89 ± 93** | **77 ± 33** | **491 ± 177** | **n.d.** | **132 ± 111** | **81 ± 100** |
|  |  | **28** | **328 ± 261** | **n.d.** | **72 ± 70** | **67 ± 32** | **814 ± 322** | **n.d.** | **127 ± 87** | **79 ± 93** |
| Red lineage<br>Chl *a* + *c* | *D. lutheri* | 5 | 332 ± 38 | n.d. | 54 ± 19 | 57 ± 21 | 501 ± 60 | n.d. | 207 ± 62 | 150 ± 10 |
|  |  | 28 | 332 ± 43 | n.d. | 30 ± 18 | 55 ± 30 | 605 ± 61 | n.d. | 216 ± 164 | 103 ± 10 |
|  | *N. gaditana* | 5 | 486 ± 71 | n.d. | 56 ± 17 | 45 ± 40 | 468 ± 129 | n.d. | 64 ± 17 | 19 ± 6 |
|  |  | 28 | 455 ± 74 | n.d. | 41 ± 17 | 31 ± 15 | 958 ± 324 | n.d. | 65 ± 19 | 19 ± 6 |
|  | *N. salina* | 5 | 576 ± 132 | n.d. | 109 ± 117 | 55 ± 68 | 465 ± 159 | n.d. | 91 ± 34 | 52 ± 68 |
|  |  | 28 | 443 ± 79 | n.d. | 47 ± 41 | 35 ± 15 | 466 ± 133 | n.d. | 71 ± 33 | 31 ± 13 |
|  | *F. pinnata* | 5 | 246 ± 61 | n.d. | 216 ± 156 | 60 ± 30 | 851 ± 252 | n.d. | 160 ± 87 | 809 ± 546 |
|  |  | 28 | 300 ± 114 | n.d. | 260 ± 178 | 75 ± 21 | 2136 ± 1369 | n.d. | 178 ± 66 | 531 ± 408 |
|  | *P. tricornutum* | 5 | 371 ± 81 | n.d. | 145 ± 50 | n.d. | 763 ± 383 | n.d. | 238 ± 68 | 82 ± 22 |
|  |  | 28 | 375 ± 75 | n.d. | 155 ± 53 | 48 ± 15 | 864 ± 463 | n.d. | 185 ± 47 | 83 ± 45 |
|  | *T. pseudonana* | 5 | 670 ± 150 | n.d. | 84 ± 22 | 114 ± 49 | 714 ± 298 | n.d. | n.d. | 45 ± 21 |
|  |  | 28 | 607 ± 90 | n.d. | 61 ± 16 | 108 ± 37 | 2812 ± 527 | 18 ± 1 | 23 ± 8 | 39 ± 18 |
|  | *T. weissflogii* | 5 | 227 ± 93 | n.d. | 53 ± 30 | 40 ± 14 | 561 ± 66 | 12 ± 1 | 116 ± 61 | 39 ± 11 |
|  |  | 28 | 183 ± 34 | n.d. | 30 ± 9 | 83 ± 98 | 644 ± 117 | n.d. | 131 ± 85 | 36 ± 27 |
|  | *P. purpureum* | 5 | 532 ± 83 | n.d. | 169 ± 123 | 66 ± 48 | 1218 ± 448 | n.d. | 243 ± 90 | 35 ± 21 |
|  |  | 28 | 482 ± 94 | n.d. | 206 ± 144 | 127 ± 170 | 1619 ± 693 | n.d. | 276 ± 225 | 49 ± 34 |
|  | *R. reticulata* | 5 | 70 ± 23 | n.d. | 118 ± 88 | 44 ± 11 | 920 ± 948 | n.d. | 110 ± 68 | 21 ± 13 |
|  |  | 28 | 58 ± 24 | n.d. | 103 ± 87 | 57 ± 28 | 1108 ± 916 | n.d. | 101 ± 51 | 27 ± 17 |
|  | *A. carterae* | 5 | 192 ± 53 | n.d. | 163 ± 266 | 106 ± 86 | 676 ± 685 | 12 ± 3 | 177 ± 143 | 101 ± 45 |
|  |  | 28 | 161 ± 48 | n.d. | 128 ± 173 | 70 ± 39 | 750 ± 696 | 9 ± 1 | 98 ± 45 | 84 ± 37 |
|  | *A. klebsii* | 5 | 203 ± 33 | n.d. | 31 ± 33 | 76 ± 14 | 417 ± 78 | 5 ± 2 | 94 ± 11 | 52 ± 8 |
|  |  | 28 | 246 ± 34 | n.d. | 20 ± 8 | 69 ± 22 | 573 ± 63 | 7 ± 1 | 108 ± 32 | 60 ± 10 |
|  | **Red lineage** | **5** | **355 ± 189** | **n.d.** | **109 ± 59** | **66 ± 25** | **687 ± 243** | **10 ± 4** | **150 ± 64** | **128 ± 229** |
|  |  | **28** | **331 ± 162** | **n.d.** | **98 ± 80** | **69 ± 29** | **1140 ± 747** | **11 ± 6** | **132 ± 74** | **97 ± 147** |
| Cyano-<br>bacteria<br>Chl *a* +<br>phyco-cyanin | *A. brevis* | 5 | 60 ± 32 | n.d. | 153 ± 84 | 31 ± 2 | 1253 ± 1184 | n.d. | 141 ± 69 | 58 ± 44 |
|  |  | 28 | 59 ± 26 | n.d. | 191 ± 173 | 39 ± 12 | 2366 ± 2108 | n.d. | 121 ± 72 | 75 ± 70 |
|  | *Synechococcus*<br>*sp.* | 5 | 100 ± 105 | 170 ± 66 | 213 ± 189 | 21 ± 3 | 436 ± 90 | n.d. | 66 ± 8 | 31 ± 49 |
|  |  | 28 | 32 ± 21 | 85 ± 26 | 98 ± 56 | 54 | 747 ± 250 | n.d. | 70 ± 8 | 47 ± 39 |
|  | **Cyanobacteria** | **5** | **80 ± 28** | **170** | **183 ± 42** | **26 ± 7** | **845 ± 578** | **n.d.** | **104 ± 53** | **45 ± 19** |
|  |  | **28** | **46 ± 19** | **85** | **145 ± 66** | **47 ± 11** | **1557 ± 1145** | **n.d.** | **96 ± 36** | **61 ± 20** |

Changes in elemental concentrations of all the species to the increased sulfate availability from 5 to 28 mM are shown in Table 3. The table reveals a correlation between changes in carbon (C) and nitrogen (N) concentrations concerning sulfate levels in the media: when C concentration decreased at 28 mM sulfate compared to 5 mM, there was a corresponding decrease in N, and vice versa, and, in addition, species that maintained stable C concentrations also exhibited no significant change in N. The concentration of P in algae of the red lineage was 17% higher at low sulfate concentration (averaging 116 mM and 97 mM at 5 and 28 mM sulfate, respectively; *p* ≤ 0.01), while algae of the green lineage contained similar P concentrations (100 and 99 mM P for 5 and 28 mM S respectively, Figure 1). Among the red lineage species, eight out of 11 had variations in P concentrations in response to different S concentrations in the media with six of them down-regulating P at 28 mM sulfate (Table 3). In contrast, among the green lineage species, there was minimal change in P concentration when grown at different S concentrations: only two out of the seven responded in opposite directions with low statistical significance (*p* ≤ 0.1) (Table 3). Surprisingly, the variance of sulfate availability did not translate to S cell concentration (Table 3). Changes were evident only in four out of 11 species from the red lineage and two out of 7 species of the green lineage with only half of them displaying higher S concentration with increased sulfate in the medium. Additionally, K concentration in both cyanobacteria species increased at 28 mM S, albeit only for *A. brevis* when calculated as cell quota. Notably, neither the red lineage nor the green lineage species showed significant fluctuation in K concentration, with response equally distributed among the lineages and devoid of a discernible trend for up or down regulation (Table 3).

**Table 3.**
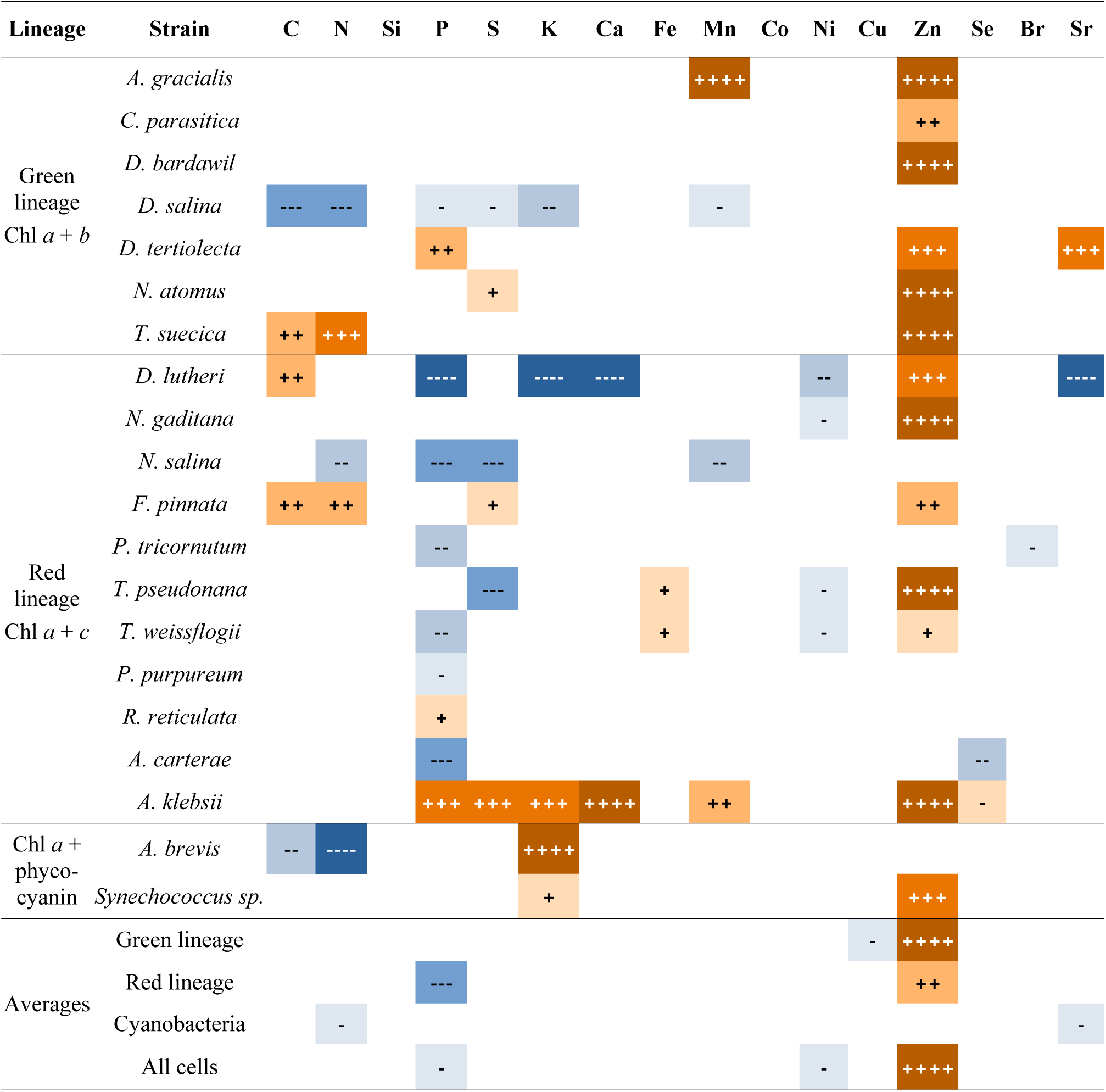
Changes in the elemental concentration in monocultures grown with different sulfate concentrations. Orange and + indicate that the concentration of the element is significantly larger in monocultures grown in high sulfate concentration (28 mM), while blue and -indicate that the concentration is significantly larger in cells grown at lower sulfate concentration (5mM). From darker to lighter shades, the probabilities are *p* ≤ 0.001(++++/−−−−), *p* ≤ 0.01(+++/−−−), *p* ≤ 0.05(+ +/−−), and *p* ≤ 0.1(+/−). Unfilled boxes represent elements that were not affected by sulfate availability.

While the influence of sulfate supply on the macronutrients examined (C, N, P, S and K) showed a clear trend only for P, a notably larger response magnitude was found for the micronutrient Zn. The majority of species cultivated with high sulfate concentration exhibited an increase in their Zn content. This up-regulation of Zn was clearly independent of genotypic constraints or plastid type (Table 3). Only seven out of 20 species did not show an increase in Zn concentration in response to elevated sulfate, and among them, five down-regulated P. Furthermore, regardless of S availability, the average Zn concentration for the red lineage algae was significantly higher than that of the algae of the green lineage (*p* ≤ 0.01, Table 2, Figure 1).

Mn cell concentration was altered by S availability in only four species (Table 3), however, no overall trend was identified related to lineage. Four species showed lower Ni at 28 mM sulfate, and none showed an opposite response (Table 3). Our results indicate that there might be a potential down-regulation of Ni in response to high sulfate in some red linage algae with low statistical significance (*p* ≤ 0.1) (Table 3). Se, both on a cell and volume basis, was affected by sulfate concentration in the medium only in two *Amphidinium* species albeit in opposite directions: *A. klebsii* had higher Se concentration at high S availability, whereas lower in *A. carterae* (Table 3). Se was reliably detected only in these two dinoflagellates, while in other species it was found in an insufficient number of replicate readings to reliably establish its presence. The concentration of both Ca and Br was less responsive to sulfate availability than when calculated in terms of cell quota (Table 3), and other trace elements, like Cu or Co, were not affected by the external sulfate concentration. Unexpectedly, we consistently obtained readings for Cr at non-negligible levels. This was surprising since no Cr was intentionally added to our media, however we cannot exclude that low levels of contaminants were present in the reagents we used to prepare the media, or in the reagents used to prepare the samples.

### Effects of S availability on elemental stoichiometry

The C:N, C:P, N:P, S:P and C:S ratios for all species studied at both low and high sulfate conditions are presented in Table 4. Figure 2 shows box plots of the same ratios grouped by lineages and cyanobacteria at both low and high sulfate conditions. As expected, the C:N ratio exhibited the most consistent stability across various lineages, species and sulfate conditions (Fig. 2). Differences in C:N grown under different sulfate concentrations were identified for both the green and red lineages, but the significance was weak (*p* = 0.04 and 0.08, respectively). In contrast, the C:P and N:P ratios demonstrated substantial variability among species (Table 4), particularly with higher values at high sulfate concentration. The mean and the variability of these ratios in the green lineage were highest (Table 4, Fig. 2). Specifically, under equivalent sulfate concentrations, the C:P and N:P ratios in the green lineage significantly surpassed those of the red lineage (*p* ≤ 0.05).

**Figure 2.**
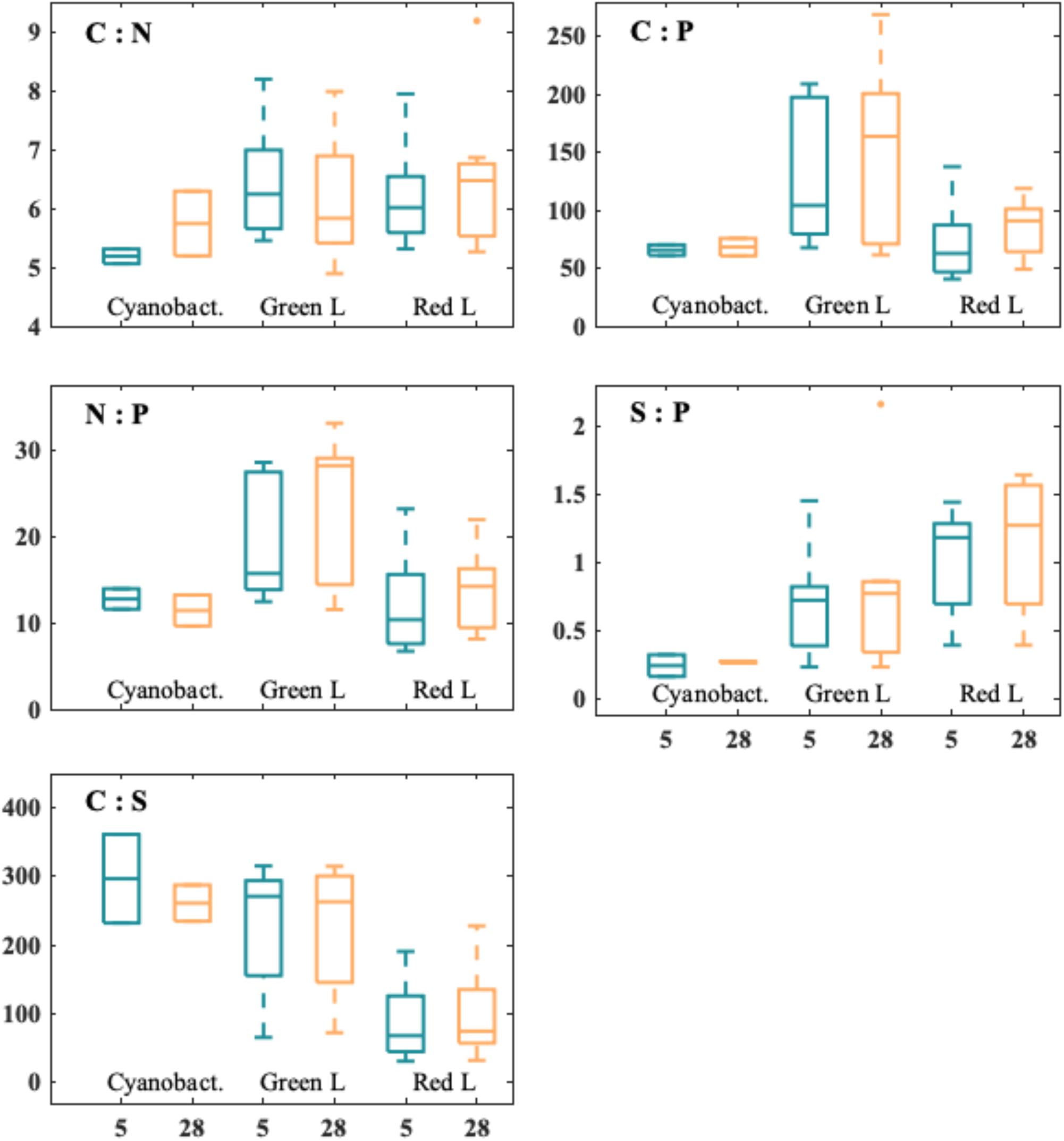
Boxplots showing changes in C:N, C:P, N:P, S:P and C:S stoichiometries of Cyanobacteria (n=2), and species from the Green (n=7) and from the Red (n=11) Lineage grown with 5 mM (orange) or 28 mM (blue) of sulfate concentration in the media. Note that y-axes change.

**Table 4.** Molar stoichiometry of C:N, C:P, N:P, S:P, and C:S (average ± standard deviation) in the studied species when grown with different sulfate availability.

| Lineage | Strain | SO<sub>4</sub><sup>-</sup> (mM) | C : N | C : P | N : P | S : P | C : S |
| --- | --- | --- | --- | --- | --- | --- | --- |
| Green<br>lineage<br>Chl *a* + *b* | *A. gracialis* | 5 | 7.2 ± 0.9 | 202.1 ± 39.2 | 28.2 ± 4.5 | 0.72 ± 0.1 | 280.1 ± 49.9 |
|  |  | 28 | 7.0 ± 0.8 | 204.6 ± 42.7 | 29.1 ± 3.9 | 0.77 ± 0.1 | 269.6 ± 57.3 |
|  | *C. parasitica* | 5 | 6.4 ± 0.2 | 184.6 ± 41.7 | 28.5 ± 5.9 | 0.84 ± 0.1 | 221.6 ± 55.9 |
|  |  | 28 | 6.6 ± 0.2 | 190.0 ± 52.5 | 28.7 ± 7.7 | 0.85 ± 0.2 | 220.8 ± 30.5 |
|  | *D. bardawil* | 5 | 8.2 ± 0.7 | 209.2 ± 137.2 | 25.1 ± 15.3 | 0.76 ± 0.1 | 271.2 ± 162.5 |
|  |  | 28 | 8.0 ± 0.8 | 269.1 ± 118.5 | 33.1 ± 12.3 | 0.86 ± 0.2 | 311.3 ± 112.2 |
|  | *D. salina* | 5 | 5.4 ± 0.8 | 67.7 ± 12.7 | 12.4 ± 1.3 | 0.23 ± 0.0 | 298.7 ± 37.4 |
|  |  | 28 | 5.4 ± 0.4 | 61.4 ± 17.5 | 11.5 ± 3.4 | 0.23 ± 0.0 | 263.3 ± 66.7 |
|  | *D. tertiolecta* | 5 | 6.2 ± 0.8 | 104.3 ± 17.9 | 15.0 ± 6.2 | 0.33 ± 0.0 | 315.6 ± 90.0 |
|  |  | 28 | 5.5 ± 0.2 | 80.4 ± 22.5 | 14.5 ± 4.1 | 0.26 ± 0.0 | 315.1 ± 87.5 |
|  | *N. atomus* | 5 | 5.5 ± 0.1 | 73.3 ± 8.3 | 13.4 ± 1.5 | 0.55 ± 0.0 | 133.9 ± 20.5 |
|  |  | 28 | 4.9 ± 1.8 | 67.9 ± 27.7 | 14.4 ± 2.6 | 0.57 ± 0.1 | 120.9 ± 47.7 |
|  | *T. suecica* | 5 | 6.2 ± 0.2 | 98.0 ± 42.2 | 15.7 ± 6.7 | 1.45 ± 0.4 | 65.8 ± 12.2 |
|  |  | 28 | 5.8 ± 0.3 | 163.9 ± 127.7 | 28.2 ± 22.0 | 2.16 ± 1.4 | 72.6 ± 10.4 |
|  | **Green lineage** | **5** | **6.5 ± 1.9** | **134.2 ± 62.1** | **19.8 ± 7.2** | **0.70 ± 0.4** | **226.7 ± 93.5** |
|  |  | **28** | **6.2 ± 1.1** | **148.2 ± 80.0** | **22.8 ± 8.9** | **0.81 ± 0.6** | **224.8 ± 94.1** |
| Red<br>lineage<br>Chl *a* + *c* | *D. lutheri* | 5 | 6.6 ± 0.2 | 44.3 ± 8.9 | 6.7 ± 1.3 | 0.42 ± 0.0 | 108.9 ± 29.2 |
|  |  | 28 | 6.8 ± 0.4 | 88.1 ± 17.7 | 13.0 ± 3.0 | 0.73 ± 0.1 | 121.2 ± 23.8 |
|  | *N. gaditana* | 5 | 6.3 ± 0.5 | 87.7 ± 24.1 | 13.9 ± 3.3 | 0.68 ± 0.2 | 132.1 ± 21.4 |
|  |  | 28 | 6.6 ± 0.5 | 94.5 ± 36.1 | 14.2 ± 4.9 | 0.67 ± 0.2 | 140.2 ± 10.6 |
|  | *N. salina* | 5 | 7.9 ± 0.5 | 71.5 ± 7.6 | 9.0 ± 1.1 | 0.39 ± 0.0 | 183.9 ± 26.5 |
|  |  | 28 | 9.2 ± 0.6 | 89.8 ± 14.5 | 9.8 ± 1.5 | 0.39 ± 0.0 | 228.3 ± 35.7 |
|  | *F. pinnata* | 5 | 6.0 ± 0.3 | 40.6 ± 11.7 | 6.8 ± 1.8 | 1.25 ± 0.3 | 32.8 ± 6.0 |
|  |  | 28 | 5.9 ± 0.3 | 49.1 ± 12.3 | 8.4 ± 2.1 | 1.57 ± 0.3 | 32.3 ± 9.4 |
|  | *P. tricornutum* | 5 | 5.5 ± 0.2 | 40.9 ± 9.1 | 7.5 ± 1.7 | 1.31 ± 0.2 | 31.2 ± 5.1 |
|  |  | 28 | 5.4 ± 0.1 | 50.4 ± 15.0 | 9.3 ± 2.6 | 1.55 ± 0.4 | 33.3 ± 8.5 |
|  | *T. pseudonana* | 5 | 6.9 ± 0.6 | 54.0 ± 19.7 | 7.7 ± 2.3 | 1.29 ± 0.5 | 42.7 ± 8.9 |
|  |  | 28 | 6.9 ± 0.7 | 56.2 ± 16.0 | 8.1 ± 1.9 | 0.99 ± 0.2 | 57.6 ± 13.6 |
|  | *T. weissflogii* | 5 | 5.4 ± 0.1 | 62.8 ± 22.6 | 11.6 ± 4.1 | 1.00 ± 0.2 | 62.3 ± 13.6 |
|  |  | 28 | 5.3 ± 0.2 | 98.1 ± 39.6 | 18.6 ± 7.4 | 1.27 ± 0.2 | 76.2 ± 24.9 |
|  | *P. purpureum* | 5 | 5.3 ± 0.2 | 86.2 ± 19.3 | 16.2 ± 3.2 | 1.26 ± 0.2 | 68.4 ± 10.3 |
|  |  | 28 | 5.4 ± 0.1 | 118.9 ± 24.0 | 21.9 ± 4.2 | 1.60 ± 0.2 | 74.9 ± 15.2 |
|  | *R. reticulata* | 5 | 6.0 ± 1.0 | 137.7 ± 34.1 | 23.2 ± 4.8 | 0.73 ± 0.1 | 191.0 ± 46.3 |
|  |  | 28 | 6.5 ± 0.7 | 105.5 ± 15.8 | 16.4 ± 2.9 | 0.68 ± 0.1 | 158.1 ± 33.7 |
|  | *A. carterae* | 5 | 6.0 ± 0.3 | 61.7 ± 15.6 | 10.3 ± 2.9 | 1.18 ± 0.1 | 51.9 ± 10.9 |
|  |  | 28 | 6.2 ± 0.3 | 90.0 ± 10.1 | 14.7 ± 1.7 | 1.64 ± 0.4 | 57.4 ± 12.6 |
|  | *A. klebsii* | 5 | 6.3 ± 0.2 | 125.3 ± 19.2 | 19.8 ± 3.1 | 1.44 ± 0.1 | 87.0 ± 10.2 |
|  |  | 28 | 6.5 ± 0.2 | 102.2 ± 20.7 | 15.7 ± 3.5 | 1.37 ± 0.1 | 74.5 ± 13.1 |
|  | **Red lineage** | **5** | **6.2 ± 0.8** | **73.9 ± 32.9** | **12.1 ± 5.6** | **1.00 ± 0.4** | **90.2 ± 57.3** |
|  |  | **28** | **6.4 ± 1.1** | **85.8 ± 23.4** | **13.7 ± 4.5** | **1.13 ± 0.5** | **95.8 ± 60.1** |
| Chl *a* +<br>phyco-<br>cyanin | *A. brevis* | 5 | 5.1 ± 0.3 | 70.2 ± 10.5 | 13.9 ± 2.4 | 0.32 ± 0.1 | 232.5 ± 77.5 |
|  |  | 28 | 6.3 ± 0.2 | 60.7 ± 16.0 | 9.6 ± 2.3 | 0.27 ± 0.1 | 235.0 ± 72.9 |
|  | *Synechococcus* sp. | 5 | 5.3 ± 0.3 | 61.1 ± 10.4 | 11.6 ± 2.2 | 0.16 ± 0.1 | 361.4 ± 118.6 |
|  |  | 28 | 5.2 ± 0.3 | 75.8 ± 26.8 | 13.2 ± 7.5 | 0.26 ± 0.1 | 287.7 ± 79.2 |
|  | **Cyanobacteria** | **5** | **5.2 ± 0.2** | **65.7 ± 6.4** | **12.7 ± 1.7** | **0.24 ± 0.1** | **296.9 ± 91.1** |
|  |  | **28** | **5.7 ± 0.8** | **68.3 ± 10.7** | **11.4 ± 2.5** | **0.26 ± 0.0** | **261.4 ± 37.3** |
| All cells | **Green lineage** |  | 6.3 ± 1.0 | 141.2 ± 69.2 | 21.3 ± 7.9 | 0.8 ± 0.5 | 225.8 ± 90.1 |
|  | **Red lineage** |  | 6.3 ± 0.9 | 79.8 ± 28.5 | 12.9 ± 5.0 | 1.1 ± 0.4 | 93.0 ± 57.4 |
|  | **Cyanobacteria** |  | 5.5 ± 0.6 | 67.0 ± 7.3 | 12.1 ± 1.9 | 0.3 ± 0.1 | 279.1 ± 60.4 |

The average C:S ratio among the 11 species belonging to the red lineage was similar at both sulfate concentrations; whereas within the green lineage it more than doubled (227 at 5 mM sulfate; 225 at 28 mM sulfate) and significantly different (*p* ≤ 0.01) from the red lineage (Fig. 2). The S:P ratio kept its homeostasis across lineages and sulfate concentration. Notably, both cyanobacteria exhibited C:S and S:P ratios more aligned with the green rather than with the red lineage (Fig. 2).

When the C:N, C:P, N:P, S:P, C:S are considered, the species belonging to the red lineage showed greater sensitivity to the change in S availability than those in the green lineage. In particular, in the red lineage, eight out of the eleven species altered at least one of the ratios to P, and among these eight species, six had higher ratios at 28 mM S (Table 4). This reflects the lower P cell quotas found in the red lineage at 28 mM sulfate compared to 5 mM sulfate (Table 3, *p* ≤ 0.01).

The Zn:P was the only trace element stoichiometry that increased when cells were grown in high sulfate concentration, in both lineages (from 5.3 ± 1.9 to 8.8 ± 3.2 in green lineage and from 7.1 ± 6.0 to 11.5 ± 6.2 in red lineage, *p* ≤ 0.001). There were no discernible differences in trace element stoichiometry between red and green lineages at the same sulfate concentration (*p* > 0.25). Ranked by their cell quota averaged over all red and green lineages, trace metal stoichiometry was as follows: Fe>Zn>Mn>Br>Ni≈Sr>Cu>Co>Se. It is worth noting that the Co value was derived from a limited dataset within cyanobacteria, included primarily for comparison with rankings from prior studies by Ho *et al*. (2003) and Twining & Baines (2013). The aggregated trace elemental stoichiometry, averaged across both sulfate conditions (note that Zn increased significantly with elevated sulfate concentration, as explained above), for the green lineage was:

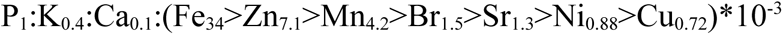

and for the red lineage was:

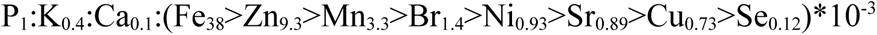

## Discussion

### Phytoplankton stoichiometry and cell quotas

Our data indicates that macronutrients stoichiometry did not change when the species studied were grown in low or high sulfate media. However, distinct differences existed in the C:P, N:P and C:S stoichiometry between lineages. The macronutrients stoichiometry averaged for all the studied species (C:N:P:S=100:16:1:0.9) followed closely the canonical Redfield ratio of 106:16:1:1.6 (Redfield *et al*., 1963), and it fell within published values (Geider & La Roche, 2002; Heldal *et al*., 2003; Martiny *et al*., 2013), however, the red lineage stoichiometry had lower values (C:N:P:S=80:13:1:1.1) than the green lineage (C:N:P:S=141:21:1:0.8, Table 4), as found by Ho *et al*. (2003). Our averaged cell P concentration, closely resembling the findings of Ho *et al*. (2003), seemingly suggests a lower absolute C and S content in the red lineage species rather than an increased P concentration. It is also therefore possible that, relative to the other macronutrients, cellular P concentration was consistently high due to our sampling at the exponential growth phase, since according to the growth rate hypothesis, low C:P ratios may reflect an increase in P-rich ribosomes to support rapid protein synthesis in fast growing cells (Elser *et al*., 2000; Sterner & Elser, 2002).

Our average S:P ratio of 0.9 is similar to reported values in the literature (Finkel *et al*., 2006), but the average S:P ratio of algae in the red lineage is larger than that of algae in the green lineage (1.06 vs. 0.76, respectively). The C:S ratio showed a marked disparity, being less than half in red than in the green lineage (about 93 vs. 226, respectively), in agreement with Ho *et al*. (2003), Norici *et al*. (2005) and Segura-Noguera *et al*. (2016). Our two cyanobacteria species had an average C:S ratio of 279, exceeding the averages of *Prochlorochoccus* and *Synechococcus* (216 and 230, respectively) reported by Heldal *et al*. (2003). Collectively, the data highlight a higher S content in the red lineage, aligning with the S facilitation hypothesis, by which red lineage algae exhibit lower S use efficiency and higher reliance on S for growth (Norici *et al*., 2005; Ratti *et al*., 2011, 2013).

The averaged stoichiometry across all species shows significantly higher level of Fe and Zn (5 and 10 times, respectively), and notably lower concentration of K (approximately 5 times) compared to the values reported by Ho *et al*. (2003), in Fe-limited species. Our high Fe cell quotas fall within a physiologically compatible range (Finkel *et al*., 2006). Diatom Fe cell content can vary by over an order of magnitude, likely due to genetic adaptations; e.g. centric diatoms may store surplus Fe (Sunda & Huntsman, 1995; Kustka *et al*., 2007). Our average K levels, both in terms of molar concentration and K:P ratio, are within the range of published values (Finkel *et al*., 2006 ; Segura-Noguera *et al*., 2016). The relatively high Zn readings in our study are also plausible, considering that Ho *et al*. (2003) report an extremely low Zn concentration in their medium, combined with a high concentration of Mn, potentially leading to an overestimation of this element (Ho *et al*., 2003). Additionally, Finkel *et al*. (2006) showed how the Zn:P ratio can be highly dependent on the intensity of light irradiation, and at our irradiance level (100µE), Zn:P ratios were over 5 times higher than at 250 µE, the irradiance used by Ho *et al*. (2003).

The average of our 20 species cultivated with different S concentrations was 120µM, standing at the high range reported by Finkel *et al*. (2006) examining only five species (20 to 120µM). Twining & Baines (2013) reported Ni:P ratios ranging from 0.2 to 8 (mmol mol^−1^), but their average ratio was close to 1, like our ratio of 1.06. In our experiments, the Ni average for all species was lower under high S conditions. Interestingly, the four species that down regulated Ni also increased Zn. Ni and Zn compete for uptake in plants (Cataldo *et al*., 1978), and a recent study showed that Ni was upregulated under Zn deficiency (Nishida *et al*., 2015). To our knowledge, we present the first direct experimental evidence of a possible relationship of Ni with S availability.

### Effect of sulfate availability on S, P and Zn cell quotas

Although our averaged S:P ratio across all species and conditions did increase under the high S concentration (*p*<0.05), rising from 0.82 to 0.93, this change remained relatively limited, given the more than 5-fold increase in S availability from the media. This is not surprising, since in S uptake and assimilation are tightly regulated by metabolic demand both in plants (Takahashi *et al*., 2011; Giordano & Raven, 2014) and in algae (Prioretti *et al*., 2014). In our study of 20 species, three exhibited increased S cell concentration at 28mM sulfate, while three showed a decrease, indicating that these were species-specific (Table 3). The marginal overall rise in the S:P ratio may at least partly be explained by the overall decrease in P concentration as S availability increased from 5 to 28mM, particularly in the red lineage (Fig. 1), contributing to the frequent increase in the element:P ratio observed in the red lineage under the high S condition, while the C:N and C:S ratios seemed less affected (Fig. 2).

To the best of our knowledge, this represents the first report of differential P cell concentrations between algae of the red and green lineages, possibly overlooked due to the common practice of using P as the benchmark for ratio calculation. This difference between lineages is evident only under conditions of 28mM S representing modern oceans, while remaining imperceptible under 5mM conditions representing ancient oceans (Fig.1), pointing to the existence of a correlation between the observed ecological success of the red lineage and its dominance in modern oceans. Moreover, our study reveals the response of Zn and P to sulfate availability: Zn content increase in the majority of species under 28mM S, and among the species that do not increase Zn, most down-regulate P. The clear up-regulation of Zn at high sulfate was evident across various genotypes or plastid types (Table 3), with no exceptions found among species. We can thus hypothesize the existence of an “either Zn or P” response to changing sulfate availability, where the P response is more characteristic in species belonging to the red lineage. Zn is a key component of hundreds of biomolecules, hence it is essential in almost every cellular metabolic process (Maret, 2012; Williams, 2012), including in phytoplankton (Sunda & Huntsman 1998; Yu & Wand, 2004). Many metallo-proteins have S in their metal binding site, including some of the major Zn-metalloproteins found in phytoplankton (Twining & Baines, 2013), such as RNA synthetase or alkaline phosphatase. Also, Zn and S are tightly linked in the protein metabolism, specifically in the biosynthesis of methionine (e.g. Takahashi *et al*., 2011) and that they have been linked in the form of zinc sulfide (ZnS) since the origins of photosynthetic life forms (Mulkidjanian & Galperin, 2009).

Cytoplasmic concentrations of free Zn^2+^ must be kept extremely low to ensure the correct metallation of enzymes and the proper function of free Zn^2+^ in signaling (Yamasaki *et al*., 2007), and so, it is subject to tight regulation (Eide, 2006). The high intracellular levels found in our experiments must be located in specific Zn reservoirs, including their binding to metallothioneins, which contain up to 30% of S-rich cystein, or partitioning to specific subcellular compartments or organelles (Vallee & Falchuk, 1993; Palmiter *et al*., 1996), most probably in acidocalcisomes (Ruiz *et al*., 2001; Lander *et al*., 2016), which are also rich in P in form of pyrophosphate and polyphosphates (Docampo & Moreno, 2001). Intracellular P reserves comprise inorganic phosphates predominantly, while nucleic acids, phospholipids, and minimally, metabolites like ATP, contribute to organic pools (Geider & La Roche, 2002). P-Zn interactions in the cell include the binding of Zn to polyphosphates (for example in acidocalcisomes), since under conditions of abundant P, the formation of polyphosphate bodies can reduce the free concentration of Zn (Perales-vela *et al*., 2006; Jensen *et al*., 1982). Conversely, Zn can sequester free phosphate causing P deficiency (Kuwabara, 1985). Increased alkaline phosphatase activity conducted by PhoA (a family of enzymes containing Zn and serving as an indicator of P deficiency) has been associated with increasing Zn concentrations (Paulsson *et al*., 2002).

To further investigate the inter-relationship between Zn and P in our results, we assembled two distinct groups based on their responses to altered sulfate availability: group A encompassed eight species that exhibited increased Zn levels without a concurrent change in P, while group B comprised five species that displayed a reduction in P without an alteration in Zn levels. Group A comprised *T. pseudonana, F pinnata, N. gaditana, N. atomus, Synechococcus sp., A. graciales, T. suecica* and *D bardawil*; and group B comprised *D. salina, P. tricornutum, P. purpurium, N. salina,* and *A. carterae*. Note that group A consists of a mix of red and green lineage species whereas in group B four out of 5 species belong to the red lineage, compatible with the idea presented above that the P response is characteristic of red algae. In group A, higher S availability could have allowed the cells to produce more metallothionines to chelate Zn, while a down-regulation of P could be the result of S-substitution of P and P-reallocation in the cell, a phenomenon observed in modern ocean sulfate levels under P-limiting conditions (Van Mooy *et al*., 2009).

For both groups, we calculated the average value of Zn, as well as the Zn:P ratio under the 28mM sulfate condition. As expected, group A had a significantly greater average Zn content than group B (1.18mM and 0.87mM, respectively, *p* ≤ 0.01), however, the Zn:P ratio between these two groups was not statistically significant. This suggests a trend toward maintaining a level of homeostasis in the element ratio, regardless of alterations in absolute Zn or P concentrations. Amid changing sulfate availability, these elements tend to “walk together” to an extent, perhaps related to the crucial roles both Zn and P play in metabolic function. According to the current literature, Zn and P quotas and dynamics have been studied mainly on Zn and P supply levels (Gao *et al.,* 2016), while our findings highlight that cellular quotas of these elements are also influenced by sulfate availability.

## Acknowledgements

This work was funded by Assemble EU FP7 research infrastructure initiative through a grant awarded to M.G. in 2013 to the project “Phytoplankton Elemental Stoichiometry and its Relation to Evolution”. We thank S. Humphries (U. of Lincoln, UK) for the English revision of the manuscript. M.S.-N. was supported by an European Union’s Horizon 2020 research and innovation programme under the Marie Sklodowska-Curie grant agreement No 661063, and by the Spanish Ministry of Science and Innovation (PTA2019-017311-I). Z.X.R. was supported by National Natural Science Foundation, China (42076206) and Guangdong Basic and Applied Basic Research Foundation, China (2020A1515011073). M.S.-N. and Z.X.R. wish to convey their sincere appreciation and admiration to the late Dr. Larisa Whitney and Prof. Mario Giordano, for their visionary dedication to the field of algal physiology research.

## Data availability

The data that supports the findings of this study are available in the main text.

## Author contributions

L.W. designed and conducted the experiments and data collection. L.W. and M.S.-N. performed statistical analyses. L.W., M. S.-N., Z.X.R., and M.G. contributed to the interpretation of the data and the discussion of the results presented in the manuscript. L.W., M. S.-N., and Z.X.R. contributed to manuscript writing. M.G. designed and supervised the work, and critically revised the first draft of the manuscript.

